# Traveling waves link human visual and frontal cortex during working memory-guided behavior

**DOI:** 10.1101/2024.03.12.584543

**Authors:** Canhuang Luo, Edward F. Ester

## Abstract

Cortical traveling waves, or global patterns of activity that extend over several centimeters of the cortical surface, are a key mechanism for guiding the spatial propagation of neural activity and computational processes across the brain. Recent studies have implicated cortical traveling waves in successful short- and long-term memory encoding, storage, and retrieval. However, human memory systems are fundamentally-action oriented: eventually, the contents of memory must be utilized to produce appropriate behaviors. Cortical traveling waves could contribute to the production and control of memory-guided behaviors by flexibly routing information between brain areas responsible for storing memory content and brain areas responsible for planning and executing actions. Here, using short-term memory as a test case, we report evidence supporting this possibility. By applying image-based analyses to published human EEG studies, we show that the initiation of a memory-guided behavior is accompanied by a low-frequency (2-6 Hz) feedforward (occipital to frontal) traveling wave that predicts intra- and inter-individual differences in response onset, while the termination of a memory-guided behavior is followed by a higher frequency (14-32 Hz) feedback (frontal-to-occipital) traveling wave. Control analyses established that neither waveform could be explained by nuisance factors (e.g., propagation of visually-evoked potentials, eye movements, or passive volume conduction). Moreover, both waveforms required an overt behavior: when participants selected task-relevant memory content and prepared but did not yet execute an action based on this content, neither waveform was observed. Our findings suggest a role for traveling waves in the generation and control of memory-guided behaviors by flexibly organizing the timing and direction of interactions between brain regions involved in memory storage and action.

---

Memory systems allow agents to use knowledge and experience gained in the past to inform behaviors in the present and immediate future. This is particularly true of working memory (WM), a capacity- and duration-limited system that forms a temporal bridge between fleeting sensory phenomena and possible actions. Human neuroimaging and EEG studies have demonstrated that the contents of WM can be decoded from patterns of activity in early visual cortex (Serences et al., 2009; Harrison & Tong, 2009; Ester et al., 2009; Sprague et al., 2014; Ester et al., 2015; Sprague et al., 2016; Wolff et al., 2017; Rademaker et al., 2019; Kwak & Curtis, 2022), and that the relative quality of stimulus-specific activation patterns are strong predictors of behavioral WM performance (Ester et al., 2013). These studies support a sensory recruitment model of WM, where WM storage utilizes the same neural machinery responsible for sensory perception. However, WM is fundamentally an action-oriented system: eventually, the contents of WM must be used to produce task-appropriate behaviors. Although recent human and animal studies have documented strong and bidirectional links between systems for WM storage and action (see, e.g., van Ede, 2020, for a recent review), little is known about how sensory brain areas implicated in WM storage communicate with frontal brain areas implicated in the selection and initiation of memory-guided behaviors.

Neural oscillations are known to coordinate communication between distal brain areas (e.g., Fries, 2023). These oscillations often take the form of traveling waves, or spatially coherent oscillations that propagate across the cortex. Traveling waves of neural activity have been identified in multiple species (e.g., rodents, non-human primates, and humans) and at multiple spatial scales (see Muller et al. 2018 for a recent comprehensive review). Experimental and theoretical studies suggest that macroscopic traveling waves – i.e., global patterns of activity that extend over several centimeters of the cortical surface – are a key mechanism for guiding the spatial propagation of neural activity and computational processes across the brain (Zhang et al., 2018; Muller et al., 2018; Xu et al., 2023). Planar, rotating, and radial macroscopic traveling waves have been documented in human electrocorticographic recordings during short- and long-term memory (Zhang et al., 2018; Mohan et al., 2024; Das et al., 2024), while rotating and planar mesoscopic (intra-areal) traveling waves have been documented in monkey prefrontal cortex during WM encoding and retrieval (Bhattacharya et al., 2022). Thus, meso- and macroscopic traveling waves represent a candidate mechanism for coordinating communication between sensory and motor areas supporting memory-guided behaviors.

To test this possibility, we re-analyzed data from two published human EEG studies that independently manipulated WM storage and response demands. Image-based analyses of EEG data revealed two planar traveling waves around the time of a WM-guided behavior: A feedforward (occipital-to-frontal) low-frequency (2-6 Hz) wave that appeared shortly after a response probe and whose peak latency predicted intra- and inter-individual differences in participants’ response times, and a later feedback (frontal-to-occipital) high-frequency (14-32 Hz) wave that appeared shortly after response termination. Control analyses demonstrated that neither wave could be explained by nuisance factors (e.g., feedforward propagation of visually evoked responses, saccadic eye movements, or passive volume conduction). Critically, both waveforms required an overt behavioral response: when participants selected task-relevant memory content and prepared but did not yet execute an action based on this content, neither the feedforward low-frequency nor the feedback high-frequency wave was observed. Collectively, these results implicate traveling waves in the generation and control of memory-guided behaviors by flexibly organizing the timing and direction of network interactions across the cortex.

## Results

### Overview

To identify cortical traveling waves during WM-guided behaviors, we re-analyzed data from two experiments in two previously published EEG studies (hereafter Experiments 1 and 2; van Ede et al., 2019a; Boettcher et al., 2021, respectively). Both experiments share a similar structure. We first introduce Experiment 1 and later return to Experiment 2. In Experiment 1, human participants (N = 25) performed a WM recall task that fused components of classic WM and motor control experiments (Figure 1A).

**Figure 1.**
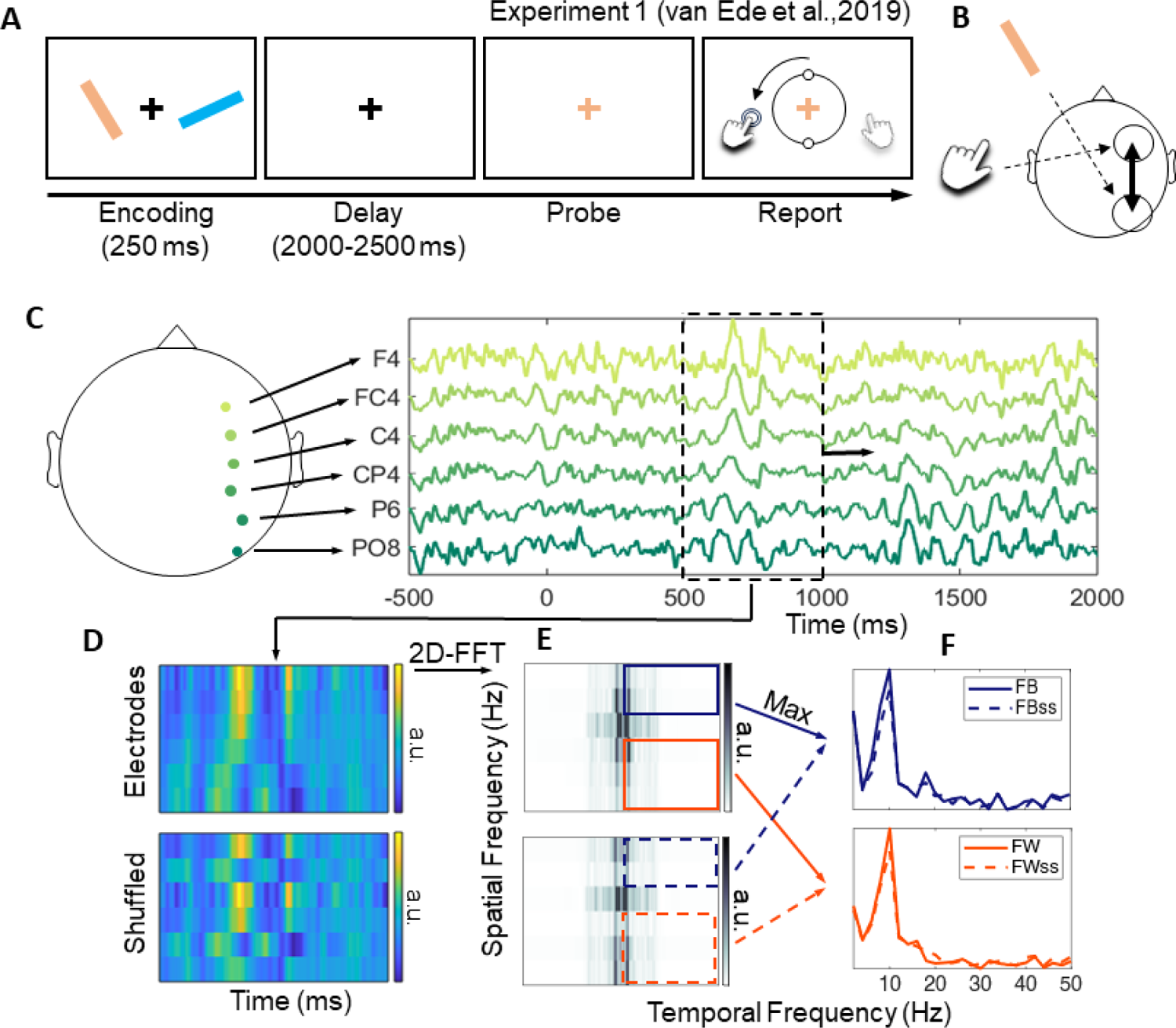
Experiment 1 Schematic and Analytic Approach. (A) Schematic of the orientation recall task used in Experiment 1 (van Ede et al., 2019a). See text for details. (B) We quantified traveling waves over planar axes contralateral to the location of the retrospectively cued response stimulus (i.e., left vs. right visual hemifield) and the task-relevant response hand (left vs. right). In this example, the storage and report of a left visual field stimulus with the left hand should require communication between right hemisphere visual and frontal areas. (C, D) We defined a planar axis over the task-relevant cerebral hemisphere and created images of EEG activity via a sliding window analysis. (E, F) We used Fourier analysis to extract frequency and phase information from each EEG image and compared forward (occipital-to-frontal; FW) and backward (frontal-to-occipital; FB) phase gradients with surrogate distributions (FWss and FBss) obtained after shuffling electrode labels. See text for additional details.

Participants remembered the orientations of two colored bars over a short delay. A subsequent change in the color of a central fixation cross (the probe) instructed participants to recall the orientation of the matching bar. Importantly, the tilt of the probe-matching bar told participants which hand should be used for recall, with counterclockwise (clockwise) tilted bars requiring a left hand (right hand) response. Participants recalled the orientation of the probed bar by pressing and holding a button on a computer keyboard with the appropriate response hand. Upon participants’ initial button press, a vertically oriented recall stimulus appeared and began to rotate in the selected direction (i.e., clockwise vs. counterclockwise). Participants were instructed to hold the response key down and release it when the orientation of the recall stimulus reached their memory of the probe-matching bar. Participants performed this task well, reporting the orientation of the probed bar with an average (±1 S.E.M.) absolute recall error of 10.69° (±0.39°) and an average post-probe response onset time of 654 (±32) ms.

A key feature of Experiment 1 (and Experiment 2) is that the physical location (i.e., left vs. right visual field) and vertical tilt (i.e., clockwise vs. counterclockwise) of the probe-matching bar were independently manipulated across trials. For example, a probe-matching stimulus initially presented in the left visual field was equally likely to require a left-vs. right-hand response. This manipulation, coupled with the contralateral organization of human sensorimotor systems, allowed us to make trial-wise predictions about the cerebral hemispheres responsible for the storage of task-relevant WM content (i.e., contralateral to the visual hemifield containing the probed bar) and the cerebral hemisphere responsible for producing the appropriate response (i.e., contralateral to the appropriate response hand). For example, we reasoned that participants’ report of a left visual field stimulus using the left hand would require coordination between right visual and right motor cortex (Figure 1B).

### Measuring Traveling Waves from Human EEG Activity

To examine whether and how cortical traveling waves contribute to WM-guided behaviors, we applied image-based analyses to EEG recordings obtained while participants performed the task summarized in Figure 1A. Since our primary goal was to examine cortical dynamics around the commission of a WM-guided behavior, we time-locked our analyses to the probe stimulus that instructed participants to recall an item held in WM. We focused our initial analyses on trials where WM storage and WM-guided responses were assumed to depend on the same cerebral hemisphere, e.g., a left (right) visual field stimulus requiring a left (right) hand response (see below for an analysis of cross-hemisphere wave propagation).

Following earlier work (Alamia et al., 2023), we defined planar axes linking occipital and frontal electrode sites over each cerebral hemisphere (Figure 1B) and constructed a series of voltage by time images (Figure 1C, D) by sliding a 500 ms window (50 ms step size) across the EEG signal measured along each axis. We applied a 2-D Fourier transform to each image, yielding a series of spectrographs where the upper and lower quadrants represent energy moving backward (i.e., frontal-to-occipital) and forward (i.e., occipital-to-frontal) along the planar axes (i.e., phase gradients; Figure 1E). We defined frequency-specific forward (FW) and backward (FB) traveling wave power by squaring the absolute value of the complex-valued Fourier coefficients in the top and bottom right quadrants of each spectrograph. We normalized FW and FB wave power by comparing observed power estimates to estimates from surrogate distributions obtained by re-computing FW and FB wave power after shuffling electrode labels 200 times, expressing total FW and FB wave power as a decibel (dB) ratio between the empirical and averaged surrogate data (Figures 1F; 2A). This step allowed us to compare FW and FB wave power across different times, experimental trials, and participants (Alamia et al., 2023; Luo et al., 2021). Finally, we compared normalized traveling wave power across cerebral hemispheres contralateral vs. ipsilateral to the location of the retrospectively probed WM item and response hand, yielding a lateralized estimate of FW and FB wave power (Figure 2B). We evaluated the statistical significance of wave power at each temporal frequency through non-parametric cluster-based randomization tests (see Methods). These analyses revealed significant FW wave power in the theta band (2-6 Hz; Figure 2B, left) and FB wave power in the beta band (14-32 Hz; Figure 2B, right). To examine the temporal dynamics of these signals, we averaged waveform power estimates within each frequency band, yielding a single FW theta and FB beta time series for each participant (Figure 3A).

**Figure 2.**
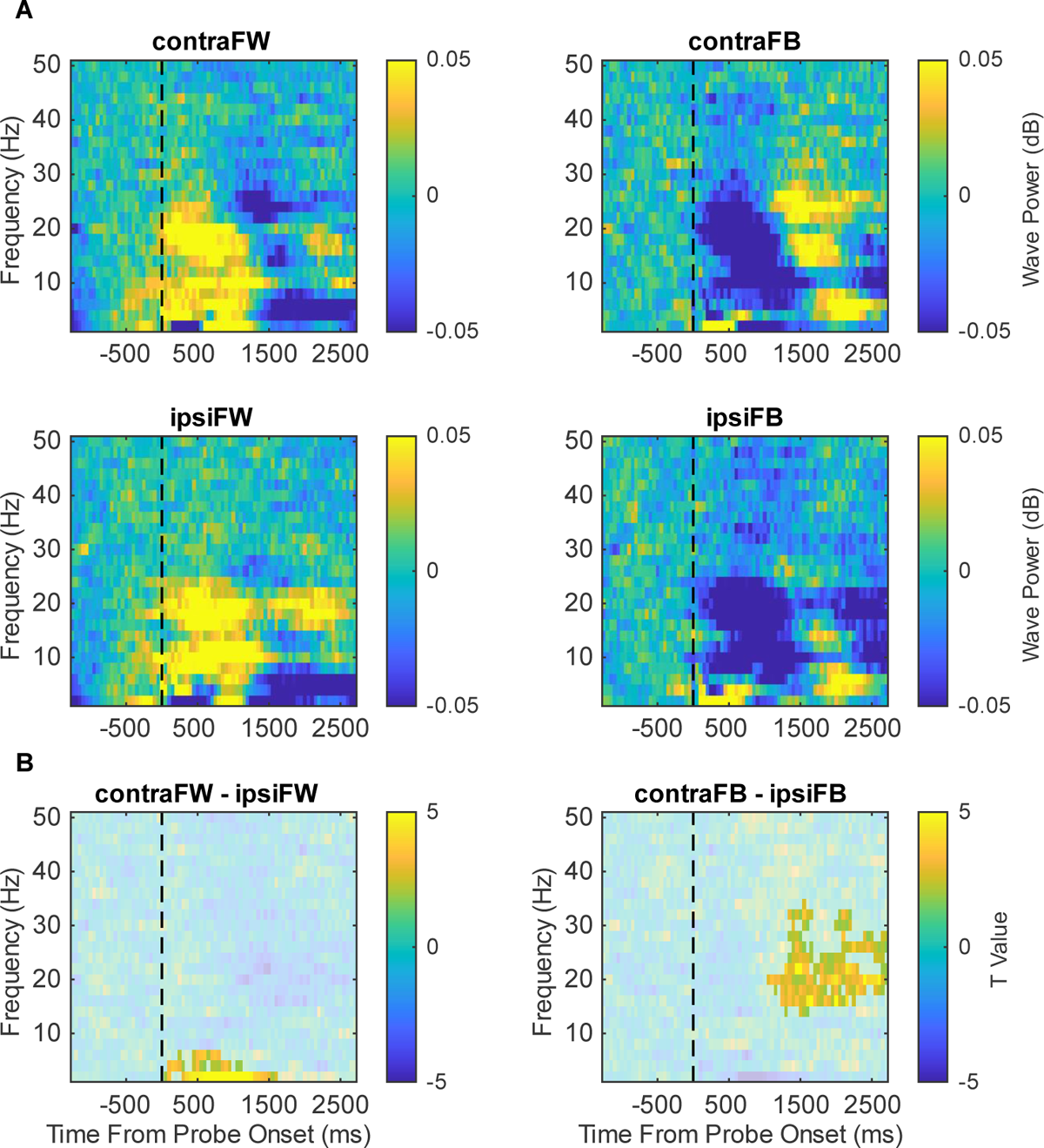
Summary of Probe-Locked Traveling Waves. (A) FW (occipital-to-frontal) and FB (frontal-to-occipital) TW power measured over the task-relevant (contralateral) and task-irrelevant (ipsilateral) cerebral hemispheres. Data in each plot have been baseline corrected from −1250 to −300 ms prior to probe onset. (B) Lateralized TW power, obtained by subtracting ipsilateral from contralateral TW power. The highlighted areas represent significant clusters (p < 0.001, cluster-based permutation test).

**Figure 3.**
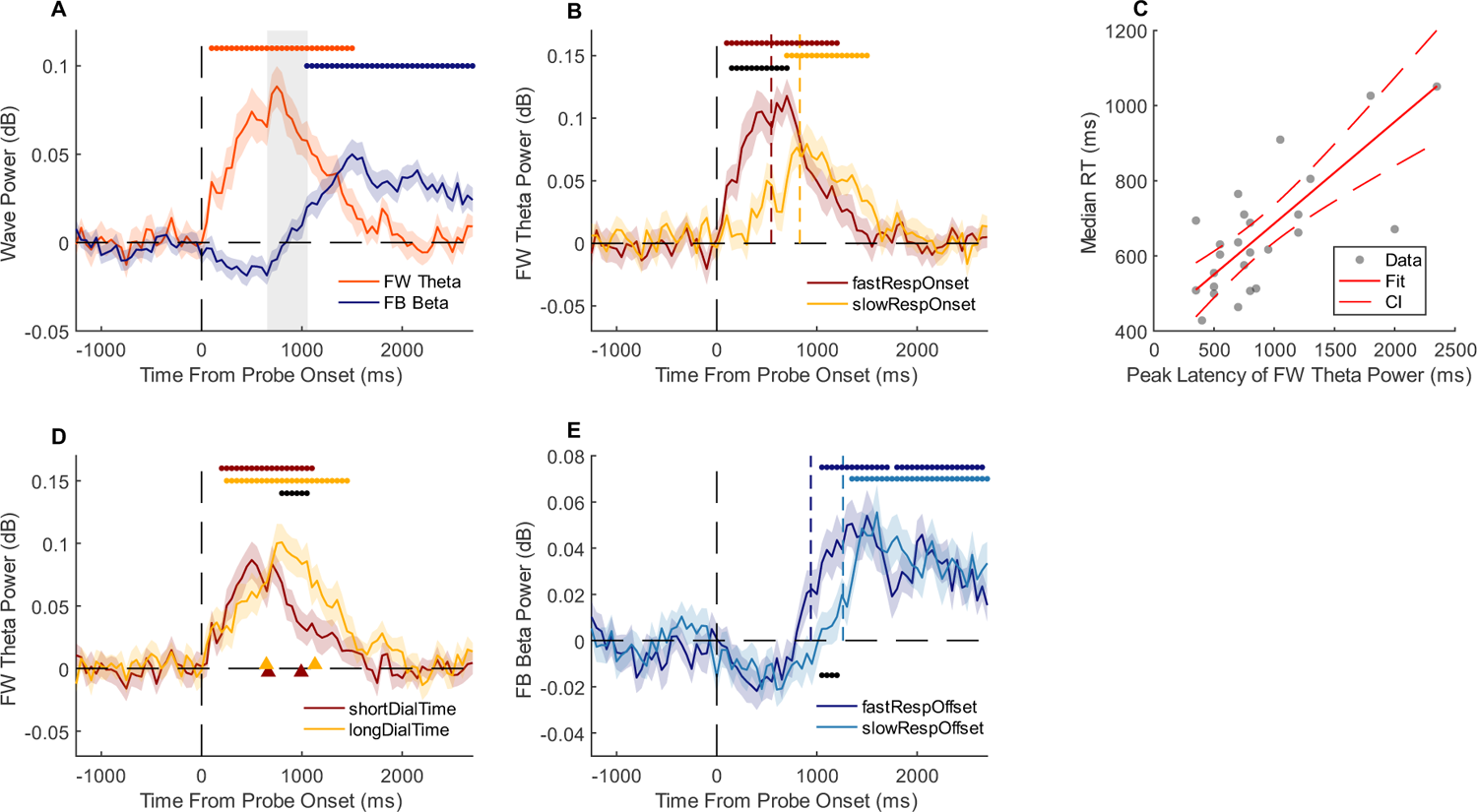
FW and FB Traveling Waves Predict Unique Aspects of Memory-Guided Behavior. (A) FW theta and FB beta wave power. The vertical dashed line at time 0 depicts probe onset; the leading and trailing edges of the grey shaded are depict median recall onset and offset across participants. (B) The timing and amplitude of FW theta wave power varied during fast response onset trials (lower 50% of all trials sorted by median response time) and slow response onset trials (upper 50%). The vertical red and yellow dashed lines depict the median response onset time in the lower and upper 50% of trials, respectively. (C) Individual differences in the peak latency of FW theta wave power predicted participants’ median response onset times across all trials. (D) FW theta wave power remained elevated throughout participants responses, lasting longer when participants spent more time rotating the recall stimulus (yellow line) compared to less time rotating the recall stimulus (red line; groups are based on a median split of total rotation time across all trials). The triangles pointing upward denote median response onset and offset corresponding to the long and short response (dial) duration trials. (E) FB beta wave power increased after participants terminated their response, and the latency of FB beta wave power increase was correlated with intra-trial variability in response offset times (i.e., the amount of time that elapsed between probe onset and the termination of participants’ responses). Shaded regions in each plot depict ±1 standard error of the mean (SEM); horizontal bars at the top and bottom of each plot depict epochs of statistical significance, with black lines denoting statistical significance between conditions (cluster-based permutation tests; p < 0.05; see Methods)

### Within-hemisphere feedforward theta and feedback beta waves predict the initiation and cessation of WM-guided behaviors, respectively

Figure 3A plots normalized FW theta and FB beta wave power time-locked to probe onset (i.e., when participants were instructed to recall the probe-matching bar). FW theta power rose quickly after probe onset and reached a maximum near participants’ median response onset time (leading edge of the vertical grey shaded area). This effect was followed by an increase in FB beta power, beginning shortly after the median response offset time (trailing edge of the vertical grey shaded area). Next, we tested these waveforms for links to behavior.

FW theta power reached a maximum near the beginning of participants’ responses (Figure 3A), consistent with a possible role in the generation and implementation of WM-guided behaviors. Under this interpretation, we made three predictions: first, the latency of FW theta power with respect to probe onset should vary with trial-wise variability in participants’ response onset times, second, differences in the average latency of FW theta power should robustly predict individual differences in response onset times, and third, the duration of increased FW theta power should vary with response duration (i.e., how long the participants held down the appropriate response button). We tested the first two predictions by quantifying FW theta power after sorting participants’ data into fast and slow response onset trials (median split; Figure 3B) and by correlating individual variability in peak theta latency (i.e., the post-probe time at which FW theta power reached its maximum) with participants’ median response onset times (Figure 3C). We tested the third prediction by quantifying FW theta power after sorting participants’ data into long and short response duration trials (where duration is defined by the difference between response onset and response offset; median split; Figure 3D). These analyses confirmed all three of our predictions: FW theta power peaked significantly earlier during fast vs. slow response onset trials (p < 0.001, cluster-based permutation test; Figure 3B), individual differences in the peak latency of FW theta power were a strong predictor of participants’ median response latencies (robust linear regression: r^2^ = 0.574, 95% bootstrap CI of r^2^ = 0.0755-0.8331; p = 6.91e-06; Figure 3C), and the duration of elevated FW theta power predicted response durations (p < 0.001, cluster-based permutation test; Figure 3D). Thus, FW theta waves are strongly linked to intra- and inter-individual differences in the initiation and duration of a WM-guided behavior. We also tested whether peak theta latency predicted intra- or individual differences in participants’ memory recall (i.e., the mean absolute distance between the correct and reported bar orientation) but found no evidence to support a link (Figure S1).

Unlike FW theta power, FB beta power reached a maximum shortly after participants terminated their response (Figure 3A). We tested FB beta waves for links with behavior by re-computing waveform power after sorting participants’ trials by response offset, that is, the total elapsed time between probe onset and participants’ final key press. Peak FB beta power was delayed by several hundred milliseconds during long-offset trials vs. short-offset trials (p < 0.05, cluster-based permutation test; Figure 3E), consistent with a mechanism operating after response termination. We did not find any evidence for a link between individual differences in peak beta waveform latency and participants’ average response offset times.

### Traveling Waves Do Not Cross Cerebral Hemispheres

Although the precise mechanisms of traveling wave conduction are unknown, both empirical and computational studies suggest a role for short-range horizontal connections between neural populations (e.g., Ermentrout & Kleinfeld, 2001; Davis et al., 2021; Sato, 2022). Conversely, communication between the cerebral hemispheres likely requires signal conduction through deeper callosal or thalamic routes with longer transduction times. Under this hypothesis, the within-hemisphere traveling waves summarized in Figures 2 and 3 may be weaker or altogether absent during trials where WM storage and response generation rely on different cerebral hemispheres (i.e., a left visual field stimulus requiring a right-hand response). We tested this prediction by computing traveling waves across axes linking left (right) occipital electrode sites with right (left) frontal electrode sites (see Methods). These analyses failed to reveal robust feedforward or feedback waveforms in any frequency band that we analyzed (2-50 Hz; Figure S2). However, note that the absence of cross-hemisphere traveling waves diminishes the possibility that the within-hemisphere waves we report are due to passive volume conduction of signals through the meninges, skull, and scalp.

### Traveling Waves are Independent of Visually Evoked Responses and Eye Movements

In control analyses, we first considered the possibility that FW theta waves were driven by visually evoked responses, i.e., the presentation of the response probe. This is unlikely: since the probe was always presented foveally it should evoke visual responses of equal magnitude across the two cerebral hemispheres, yet FW theta activity was stronger over the cerebral hemisphere contralateral to the task-relevant stimulus and appropriate response hand (Figures 2 and 3A). Nevertheless, to provide a stronger test of the hypothesis that FW theta waves reflect the feedforward propagation of a visually evoked response we re-computed FW theta power after subtracting out EEG activity phase-locked to probe onset (i.e., the visually evoked potential). This step had virtually no effect on the timing or amplitude of FW theta power (or FB beta power (Figure S3); therefore, FW theta waves are unlikely to reflect the feedforward propagation of stimulus-evoked activity.

We also considered the possibility that FW theta and FB beta waves were produced or influenced by saccadic eye movements. Large eye movements are known to produce mesoscopic traveling waves in primate visual cortex (e.g., Zanos et al., 2015) and macroscopic traveling waves in scalp EEG recordings (Giannini et al., 2018). Additionally, eye position recordings obtained during Experiment 1 indicated that participants’ gaze was subtly biased towards the physical location of the task relevant stimulus during recall (reported separately in van Ede et al., 2019b, their Experiment 1). Therefore, we tested whether traveling waves were influenced by gaze position by recomputing FW theta and FB beta waves after sorting trials according to whether they contained (a) a horizontal eye movement of any magnitude towards the location of the probed stimulus, (b) a horizontal eye movement of any magnitude towards the location of the task-irrelevant stimulus, and (c) no detectable horizontal eye movement. Trials were classified into one of these bins if horizontal gaze position exceeded a threshold of 0.057° from fixation at any time from probe onset to response termination. Although FW theta and FB beta waves were noisier (due to splitting the data into three different bins), robust theta and beta peaks were observed in each of these three bins (Figure S4). Thus, FW theta and FB beta waves are unlikely to reflect differences in eye movement patterns across trials.

### Traveling Waves Require an Overt Action

Although FW theta and FW beta waves peaked with response initiation and cessation (respectively; Figure 3A), they need not be linked with action per se. For example, these waves could reflect planning or the generation of a “tas set” that describes a plan for upcoming action (e.g., “rotate the recall stimulus by 37 degrees using your right hand” rather than the execution of that action. We tested this possibility in Experiment 2.

In Experiment 2 (Boettcher et al., 2021; their Experiment 1), human volunteers (N = 26) performed another task that fused elements of WM and motor control. Experiment 2 was identical to Experiment 1 with one exception: on many trials, a precue display presented prior to the encoding display informed participants which bar would be probed for recall with perfect accuracy (Figure 4A). Therefore, participants could select the task-relevant stimulus and associated response plan (i.e., left vs. right hand) upon presentation of the encoding display, but could not execute their response for several seconds. Indeed, spectral analyses of contemporaneous EEG recordings time-locked to the encoding display revealed decreased alpha lateralization over occipital electrode sites contralateral to the pre-cued stimulus and decreased beta lateralization over frontocentral electrode sites contralateral to the task-relevant response hand, indicating that participants selected the task-relevant stimulus and the task-appropriate response plan (Boettcher et al., 2021; their Figures 2A-C and 3A). However, although participants could (and did) select task-relevant visuomotor information from the encoding display, they were not allowed to act on this information for several seconds. This arrangement provides an ideal scenario to test whether FW theta and FB beta waves reflect the generation of a task set or an action plan. If FW theta and FB beta waves reflect the selection or preparation of a task set, then they should emerge shortly after the appearance of the encoding display and persist through the subsequent delay period.

**Figure 4.**
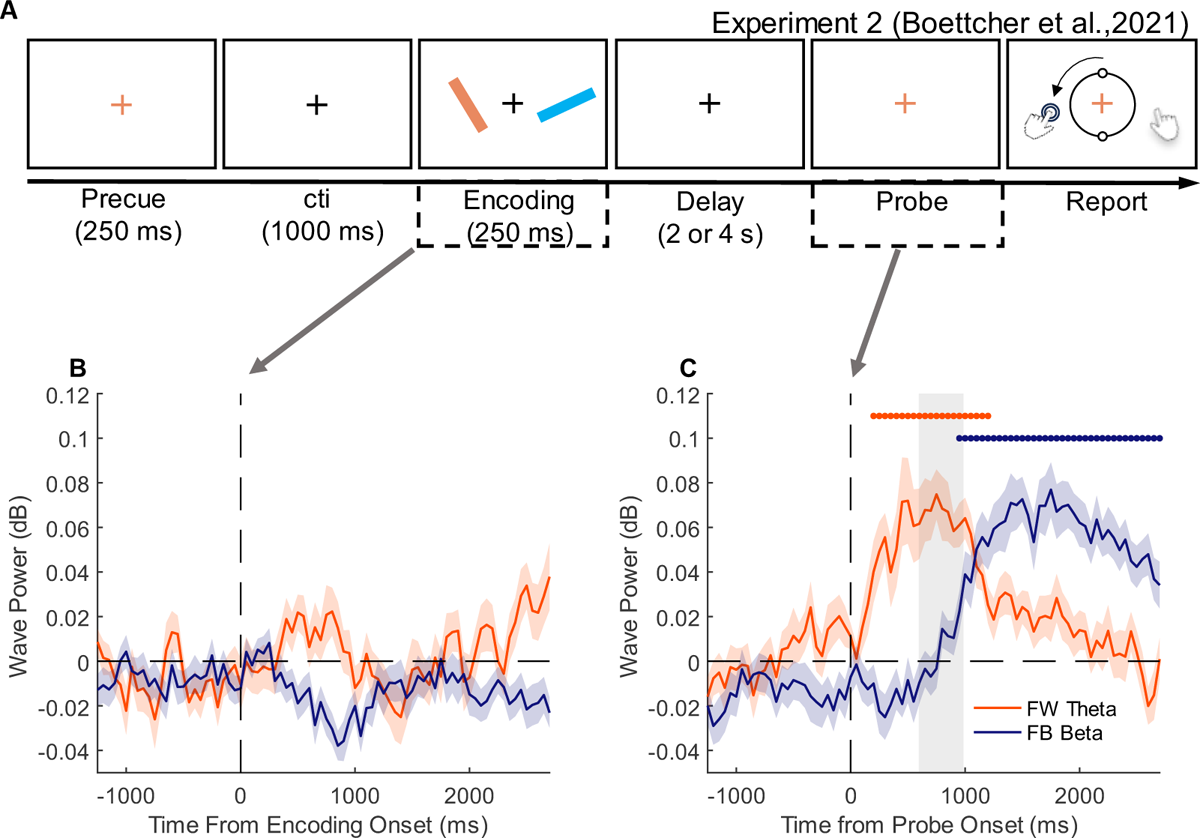
Summary of Experiment 2. (A) Experiment 2 was like Experiment 1 (Figure 1A), with the exception that the encoding display was preceded by a perfectly informative pre-cue. (B) Analysis of encoding-locked EEG data revealed neither a FW theta wave nor a FB beta wave. (C) Analysis of probe-locked EEG data revealed a clear FW theta and FB beta wave, consistent with the results of Experiment 1 (Figure 3A). Shaded regions in each plot depict ±1 standard error of the mean.

Alternately, if FW theta and FB beta waves are related to overt WM-guided behaviors, then they should only emerge after the appearance of the probe display instructing participants to initiate a response.

Using the same procedures as Experiment 1, we estimated FW theta and FB beta power time-locked to the onset of the encoding display and the onset of the probe display. Analyses time-locked to the probe display revealed a robust FW theta wave that peaked near response initiation and a FB beta wave that peaked after response termination, replicating key findings from Experiment 1 (Figure 4C; p < 0.001, cluster-based permutation test). However, FW theta and FB beta waves were absent when we time-locked our analyses to the onset of the encoding display (Figure 4B). Importantly, the lack of an effect cannot be explained by participants failing to select the task-relevant bar or task-relevant response plan following the encoding display: as described above, analyses of lateralized EEG signals time-locked to the encoding display revealed clear evidence for visuomotor selection (Boettcher et al., 2021; their Figures 2A-B and 3A). These findings support the view that FW theta and FB beta traveling waves reflect mechanisms related to overt WM-guided behaviors rather than mechanisms related to cognitive control or response preparation.

Unlike Experiment 1, we did not find any significant links between intra- and inter-individual differences in the timing of FW theta and F beta waveforms and participants’ responses (Figure S5). We suspected that the absence of such links stems from the inclusion of a perfectly informative precue, which allowed participants to select and prepare their responses prior to probe onset. Fortunately, Boettcher et al. (2021) included trials that did not contain an informative precue in their experiment. Although the number of no-precue trials was substantially less than the number of precue trials (n = ∼79 vs. ∼312), it allowed us to implement a provisional test of our hypothesis. We observed a robust correlation between individual differences in peak FW theta latency and response onset times during no-precue trials (robust linear regression: r^2^ = 0.136, p = 0.0391; Figure S6B), replicating the relationship seen in Experiment 1 (Figure 3C).

## Discussion

We examined the roles of cortical traveling waves in the coordination of WM-guided behaviors through re-analysis of two published EEG studies (van Ede et al., 2019a; Boettcher et al., 2021). Notably, both studies independently manipulated WM storage and response demands in a way that allowed us to make trial-wise predictions about the cerebral hemisphere responsible for WM storage and the cerebral hemisphere responsible for producing task-relevant responses. We identified a feedforward (posterior-to-anterior) theta wave that peaked near the initiation of a WM-guided action and that predicted intra- and inter-individual variability in response times, and a feedback (anterior-to-posterior) beta wave that peaked following the termination of a WM-guided behavior. Neither waveform could be explained by nuisance factors (e.g., feedforward propagation of stimulus-evoked responses or saccadic eye movements), and both waveforms required the generation of an overt motor response: when participants could (and did) select task-relevant WM content and linked motor plans but could not execute an overt movement for several seconds, neither waveform was observed. These observations provide important new evidence that cortical traveling waves contribute to the generation and coordination of memory-guided behaviors.

### Possible Roles for FW Theta and FB Beta Traveling Waves in WM-guided Behaviors

The precise functional roles of FW theta and FB beta waves cannot be ascertained from the data reported here. Nevertheless, the timing and direction of these waveforms allow for some reasoned speculation. FW theta waves emerged shortly after a response probe, peaked near response initiation, and remained elevated until response termination. We suggest two (non-exclusive) possibilities: first, FW theta waves could reflect a sustained channel that allows visual brain areas responsible for storing WM content to communicate with frontal areas responsible for response planning and execution (i.e., “communication through coherence”; Fries, 3. Second, FW theta waves could reflect a mechanism that compares stored memory content with external stimuli, e.g., the rotating stimulus participants adjusted to recall the target orientation. Additional experiments will be needed to test these possibilities.

FB beta waves emerged shortly after the termination of a WM-guided behavior, presumably when participants were removing now-irrelevant content from WM in anticipation of the next experimental trial. We speculate that these waveforms reflect a top-down signal that resets WM storage mechanisms in sensory cortical areas. Our speculation is based on several independent lines of evidence supporting the prediction that beta oscillations reflect top-down control over both cognitive and motor processes (see Wessel & Anderson, 2024 and Lundqvist & Miller, 2024, for recent reviews). Of particular relevance, invasive electrophysiological recordings in non-human primates suggest that short bursts of high-power beta activity enable the updating of WM contents or the removal of irrelevant information from WM (Lundqvist et al., 2018; Miller et al., 2018; Schmidt et al., 2019). Likewise, a separate human magnetoencephalography (MEG) study in patients diagnosed with obsessive compulsive disorder revealed diminished beta power that correlated with impairments in the ability to remove information from WM (Boo et al., 2023). These findings point to a role for beta oscillatory activity in updating the contents of WM or removing irrelevant information from WM. Indeed, the feedback beta waves we report were largest after the termination of a WM-guided response, presumably when participants were “resetting” WM for the next trial.

### Limitations

Scalp EEG measures cortical oscillations coherent at a spatial scale of centimeters and with geometries that encourage superposition of potentials generated by many local sources (Nunez & Srinivasan, 2006a). Signal distortion caused by volume condition makes it challenging to link EEG potentials with neocortical dynamics.

However, theoretical studies suggest that large-scale neocortical electrical fields are sufficient to produce standing and traveling oscillations that can be measured at the scalp (Nunez & Srinivasan, 2006b). In these models, local cell assemblies that drive behavior are embedded within synaptic action fields that reflect short-time changes in the number densities of active excitatory and inhibitory synapses. Recent experimental and theoretical studies suggest that electric fields carry information about WM content (Pinotsis & Miller, 2022) and organize the storage of information across multiple brain areas (Pinotsis et al., 2023). We therefore speculate that the directional and frequency-specific traveling waves reported here reflect the operation of large-scale electric fields that shape local cortical activity and information transfer across the neocortex. Further experiments and/or simulations will be needed to test this hypothesis.

The present study focused on traveling waves propagating along planar axes linking posterior and anterior electrode sites. More complicated waveforms may also contribute to WM-guided behaviors. Invasive electrophysiological studies have documented a mixture of planar and rotational (or spiral) waves in monkey prefrontal cortex during WM (Bhattacharya et al., 2022). Additionally, human iEEG studies have documented planar, rotating, and radial traveling waves that predict successful memory encoding and recall during spatial and verbal memory tasks (e.g., Zhang et al., 2018; Mohan et al., 2024; Das et al., 2024). Future research could use data-driven tools to examine whether spiral and radial waves are also present in scalp EEG data (e.g., Townsend & Gong, 2018) and explore whether these waveforms are correlated with successful short- and long-term memory performance.

The present study also focused exclusively on task-aligned traveling waves, i.e., time-locked to probe instructing participants to perform a WM-guided behavior. Spontaneous (i.e., task-independent) traveling waves are also known to contribute to visual task performance. For example, invasive electrophysiological recordings in monkey area MT have revealed the existence of spontaneous traveling waves that predict both the magnitude of target-evoked activity and target detection performance (Davis et al., 2020). Future work could examine the role(s) of spontaneous travelling oscillations in WM-guided behavior and other aspects of WM performance (e.g., encoding or storage).

### Broader Implications

Although the present study focused on WM, cortical traveling waves could reflect a general-purpose mechanism whereby sensory or mnemonic representations are mapped onto motor systems responsible for producing task-relevant behaviors. For example, human electrocorticographic studies have identified planar, spiral, and radial traveling waves during episodic long-term memory encoding and retrieval (e.g., Zhang et al., 2018; Mohan et al., 2024). Our findings complement and expand on prior work by explicitly considering how brain areas responsible for the trial-wise representation of task-relevant information in WM communicate with brain areas responsible for the trial-wise production of task-appropriate behaviors. Thus, our observations provide researchers with a new window to explore putative links between cognitive and motor control, which, with few exceptions (see van Ede et al., 2020), have heretofore been studied independently.

While the analysis and interpretation of traveling waves from scalp EEG (sEEG) activity can be challenging (see above), it does have some advantages over more invasive techniques. First, sEEG is well-tolerated by neurotypical groups across the lifespan (e.g., developmental and geriatric populations), and well-tolerated by a variety of neurodivergent groups (e.g., individuals diagnosed with socio-developmental, neurological, or psychiatric impairments). Second, sEEG emphasizes large-scale (centimeters-or-more) neocortical dynamics implicated in the control of flexible behavior (e.g., Higley & Cardin, 2022). Thus, sEEG is an excellent tool for understanding neocortical dynamics in human volunteers when medical and ethical considerations preclude invasive recordings (Lopes da Silva, 2013). Our observations give researchers and clinicians interested in understanding links between memory and action a new window with which to understand this process.

## Methods

### Overview

We reanalyzed data from two published human EEG studies. Experiment 1 used EEG data from van Ede et al. (2019) and eye tracking data from van Ede et al. (2019b); Experiment 2 used EEG data from Boettcher et al. (2021). Detailed methods can be found in each manuscript; for brevity we focus on our own methodological and analytic solutions.

### Data Availability

Raw EEG and eye tracking data for each experiment are freely available through repositories maintained by the original authors (see STAR Methods table). We are indebted to the authors for sharing their data with other researchers.

### Code Availability

Analyses reported in the current manuscript were carried out with MATLAB (Natick, MA) using open-source toolboxes (EEGLAB, Delorme & Makeig, 2004; FieldTrip, Oostenveld et al., 2011) and custom software. Analytic software sufficient to produce each manuscript figure will be made freely available upon publication of this manuscript.

### Participants

25 human volunteers (14 females) aged 19-36 completed Experiment 1. A different set of 26 human volunteers (19 females) aged 19-25 completed Experiment 2. One participant was excluded from Experiment 2 due to noncompliance with experimenter instructions. Additional details can be found in van Ede et al. (2019a) and Boettcher et al (2021). See also Table 1.

**Table 1.** Dataset Information. Trial counts refer to congruent trials (i.e., where memory and motor responses were presumed to rely on the same cerebral hemisphere).

| Study | van Ede et al, 2019a | Boettcher et al., 2021<br>Precue Trials | Boettcher et al., 2021<br>No Precue Trials |
| --- | --- | --- | --- |
| Mean ($\pm 1$ SD)<br>Trial Count (per<br>participant) | 559.8 ($\pm 25.07$) | 312.56 ($\pm 9.37$) | 78.64 ($\pm 1.05$) |
| Participant<br>Information | N = 25<br>(ages 19-36, 14 females) | N = 26<br>(ages 18-25, 19<br>females) | N = 26<br>(ages 18-25, 19<br>females) |
| Data link | https://datadryad.org/stash/dataset/doi:10.5061/dryad.sk8rb66 | https://zenodo.org/records/4471943 | https://zenodo.org/records/4471943 |

### EEG Preprocessing

Raw EEG data were high-pass filtered at 1 Hz and re-referenced to the algebraic mean of external electrodes placed over the left and right mastoids. Re-referenced EEG signals were re-sampled to 300 Hz and epoched from −1500 to +3000 ms around events of interest (e.g., onset of the probe or encoding display).

### Estimation and Quantification of Traveling Waves from EEG Data

We estimated and quantified traveling waves using a procedure developed by Almaia and colleagues (Almaia & VanRullen, 2019; Almaia et al., 2023). Trials with unusually fast or slow response onsets (>4 standard deviations from participants’ median response time) and trials where participants responded using the wrong hand were excluded from further analyses. This resulted in an average loss of 6.7% of all trials in Experiment 1 and 2.3% of trials from Experiment 2.

We defined planar axes linking frontal and electrode sites over each cerebral hemisphere (10-10 sites O1, PO3, P3, CP3, C3, FC3, and F3 for the left hemisphere and O2, PO4, P4, CP4, C4, FC4, and F4 for the right hemisphere). We generated a series of voltage-by-time images by sliding a 500 ms window (50 ms step size) across each axis (Figure 1D-E), then applied a 2-D Fourier transform to each image. The upper and lower quadrants of the resulting spectrographs depict frequency-specific phase gradients (i.e., traveling waves) moving backward (i.e., frontal-to-occipital) and forward (i.e., occipital-to frontal) along each planar axis, respectively. The upper and lower quadrants representing backward and forward phase gradients are unsymmetrical because 2D-FFT was applied on an even number of electrodes, which led to the DC component (0 on the y-axis) being on the upper quadrant. We compared observed phase gradients with surrogate distributions obtained by re-computing phase gradients after shuffling electrode labels 200 times. We expressed total wave power as a decibel (dB) ratio between the observed and averaged surrogate distributions, i.e.:

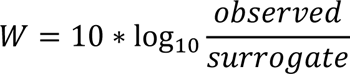

Wave power was estimated for each planar axis on each trial. Power estimates were sorted according to trial variables (i.e., the visual hemifield containing the to-be-recalled stimulus and the required response hand). For within-hemisphere analyses (Figures 2-, we subtracted wave power over the planar axis ipsilateral to the stimulus’ position and response hand from wave power over the planar axis contralateral to the stimulus’ position and response hand. Difference waves (i.e., contralateral minus ipsilateral) were then averaged across trials. Statistical significance was estimated using two-tailed non-parametric cluster-based permutation tests (Maris & Oostenveld, 2007) with a cluster forming significance threshold of 0.05.. Specifically, we compared empirically estimated difference waves with null distributions of difference waves obtained by randomly flipping the sign of each participant’s data, times. The cluster-based permutation test revealed two significant clusters in the difference waves — theta waves in the 2-6 Hz range and beta waves in the 14-32 Hz range, which were used for further analyses in Figures 2-4.

We used an identical approach to examine the propagation of traveling waves across cerebral hemispheres (Figure S1), with the exception that planar axes were defined using 10-10 electrode sites PO7, PO3, P1, CPz, C2, and FC4 (left occipital to right frontal sites) and PO8, PO4, P2, CPz, C1, and FC3 (right occipital to left frontal sites).

### Eye Position Analysis (Experiment 1)

During Experiment 1, high-fidelity gaze position estimates were obtained via an SR Research Eyelink 1000 Plus infrared eye tracking system. Binocular recordings were obtained at 1 kHz and later re-sampled to 300 Hz for alignment with contemporaneous EEG recordings. Gaze position data were used to sort experimental trials into three bins: (a) those containing a horizontal eye movement of any magnitude towards the position of the to-be-recalled stimulus, (b) those containing a horizontal eye movement of any magnitude towards the position of the task-irrelevant (i.e., non-recalled) stimulus, and (c) those containing no detectable horizontal eye movement. Trials were assigned to groups (a) or (b) if horizontal gaze position exceeded 0.057° from fixation at any point from probe onset to response termination.

## Acknowledgments

This work was supported by a grant from the National Science Foundation (NSF2050833) to EFE. We thank the authors of the original studies re-analyzed here (van Ede et al., 2019; Boettcher et al., 2021) for making their data publicly available. We would also like to thank Dr. Freek van Ede for helpful discussion and feedback on an earlier version of this manuscript.

