## Supplementary information for "Traveling waves link human visual and frontal cortex during working memory-guided behavior"

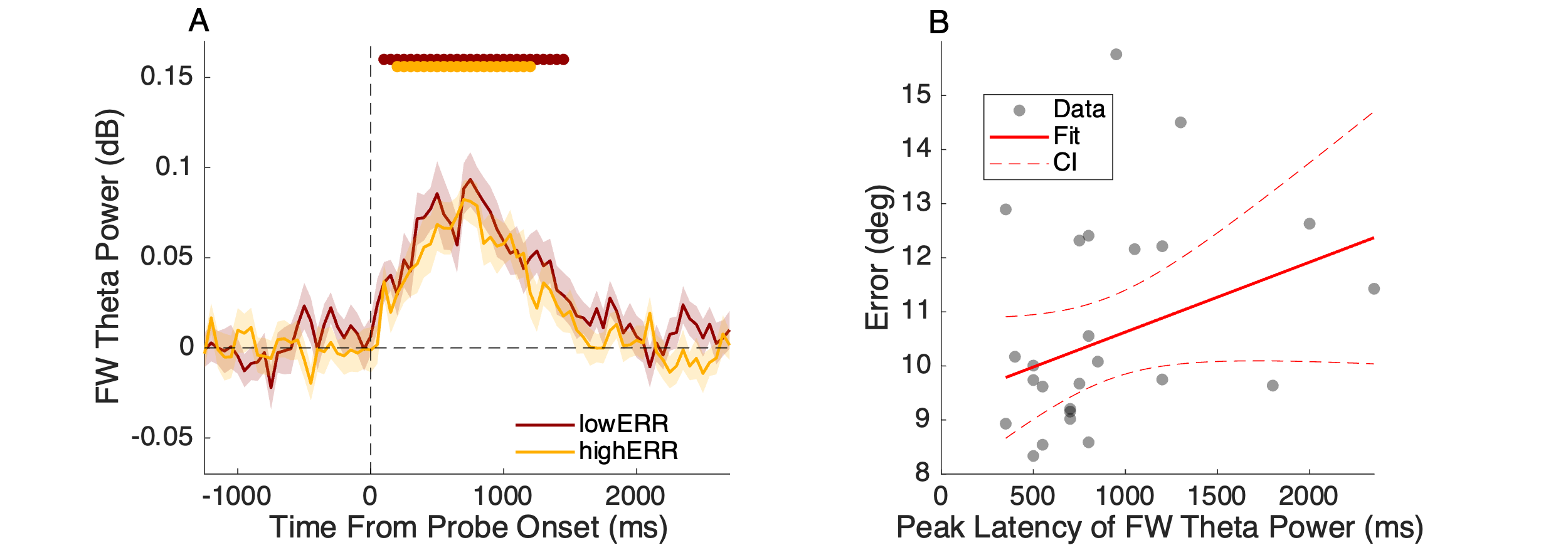

***Figure S1.*** ***FW theta latency is not correlated with response error.*** *(A) FW theta waves during low error and* *high error trials. Low and high trials were defined by applying a median split to participants average absolute recall errors. (B) Scatterplot showing no relationship between individual differences in the timing of FW theta peak amplitude and participants’ average response error (r^2^ = 0.091). Shaded regions in (A) depict the ±1 standard error of the mean; horizontal bars at the top of the plot depict epochs where wave power was significantly greater than zero (cluster-based permutation test, p < 0.05).*

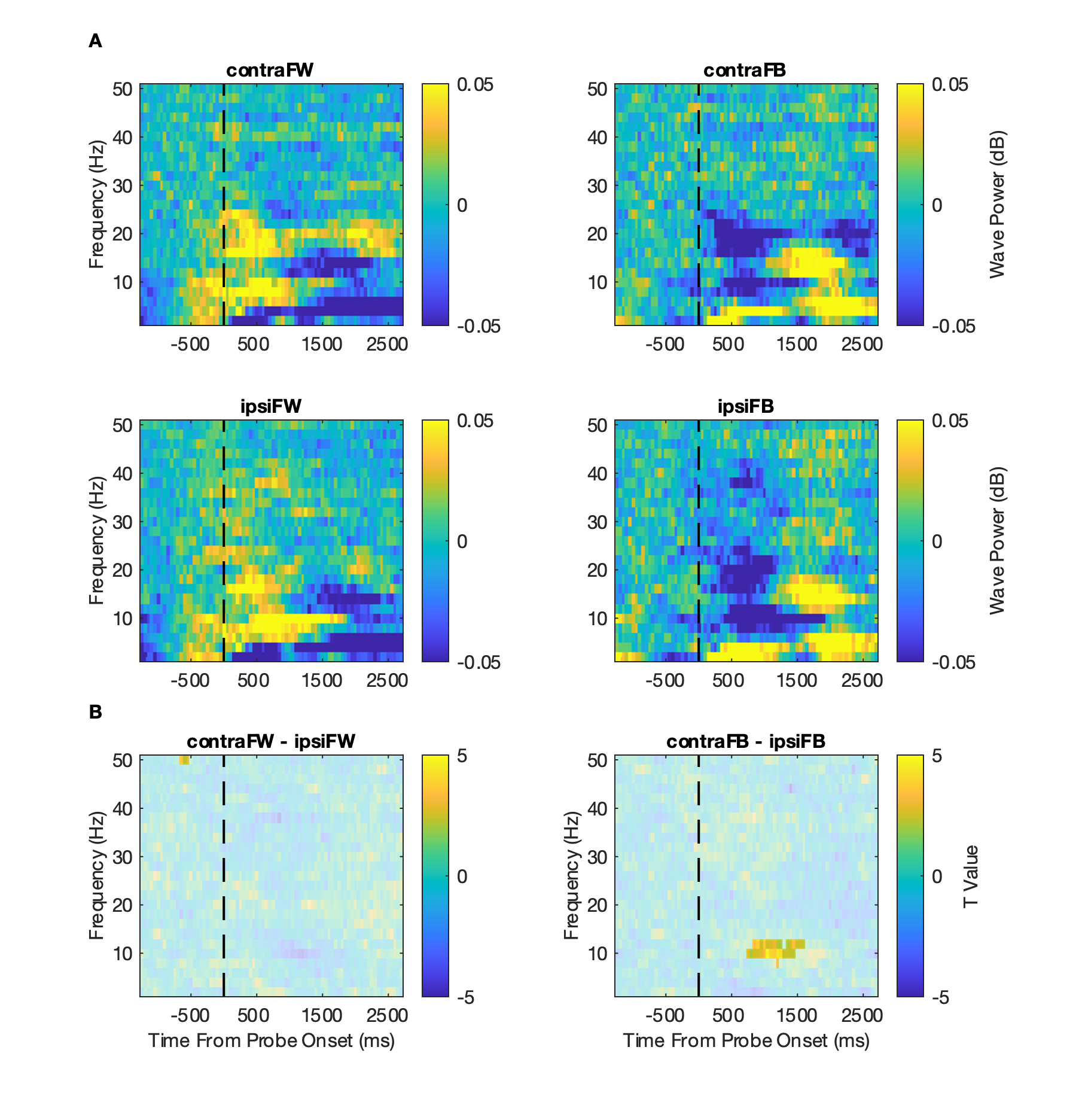

***Figure S2. TWs do not propagate across cerebral hemispheres.*** (A) Quantification of FW (left column) and FB (right column) traveling wave power across cerebral hemispheres during trials where WM storage was assumed to rely on different cerebral hemispheres (e.g., a left visual field stimulus requiring a right-hand response). Data were analyzed along (approximately) planar axes using 10-10 electrode sites PO7, PO3, P1, CPz, C2, and FC4 (left occipital to right frontal sites) and PO8, PO4, P2, CPz, C1, and FC3 (right occipital to left frontal sites).

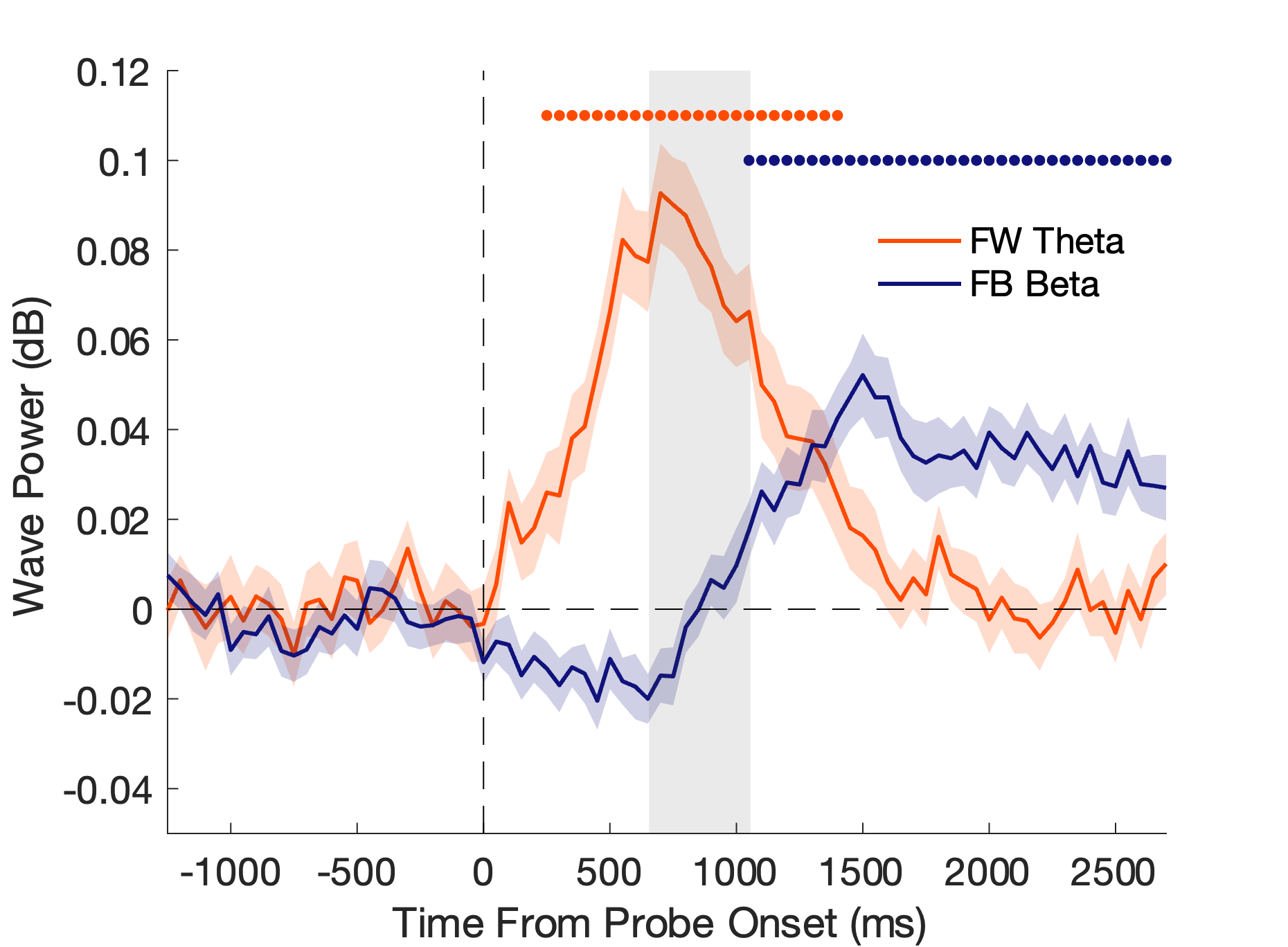

***Figure S3. FW theta and FB beta wave power after removing activity phase-locked to probe onset (i.e., the visually-evoked potential)****.* *Shaded regions around each line depict ±1 standard error of the mean; the grey shaded area depicts the median response duration (i.e., median response onset to median response offset). Horizontal bars at the top of each plot mark epochs where wave power was significantly greater than zero (p < 0.05; cluster-based permutation test).*

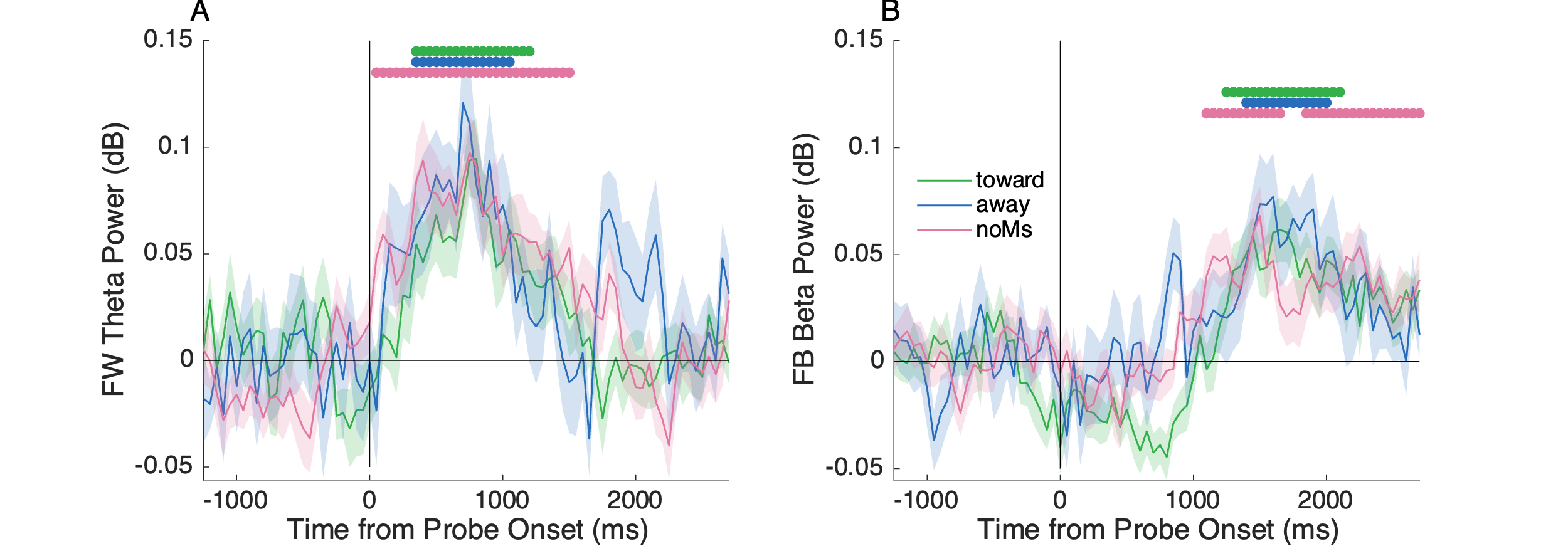

***Figure S4.*** ***FW theta and FB beta are not influenced by eye movements****. (A) FW theta wave power averaged over trials with horizontal eye movement towards the probe-matching stimulus, away from the probe-matching stimulus, and trials containing no detectable horizontal eye movement (using a threshold of 0.057° from fixation). (B) Identical to (A), but for FB beta wave power. Shaded regions in each plot depict the ±1 standard error of the mean. Horizontal bars in each plot depict epochs where wave power was significantly greater than zero (p < 0.05, cluster-based permutation test).*

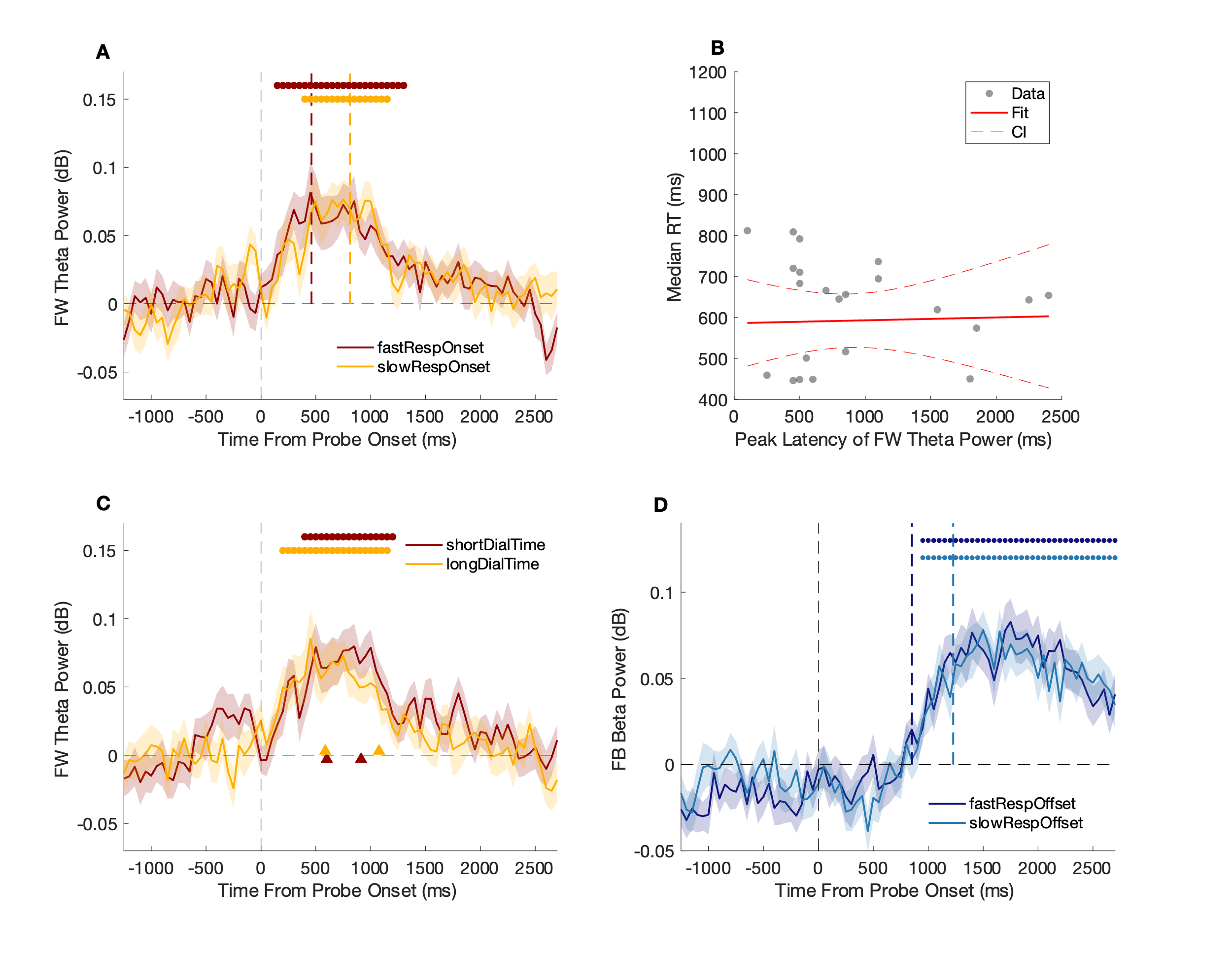

***Figure S5. No links between wave power and response times during precue trials in Experiment 2.*** *(A) Neither the amplitude nor the latency of FW theta waves varied with participants’ response onset times (median split). (B) Peak FW theta latency did not predict individual differences in response onset times. (C) Neither the amplitude nor the latency of FW theta waves varied with participants’ response duration (dial time; median split). Neither the amplitude nor the latency of FB beta waves varied with response offset times (median split). Shaded regions in each plot depict ±1 S.E.M. The vertical dashed line at time 0 in (A), (C), and (D) depict probe onset. Colored vertical dashed lines in (A), (C), and (D) depict median response onset times (fast vs. slow), response durations (short vs. long), and response offset times (fast vs. slow).*

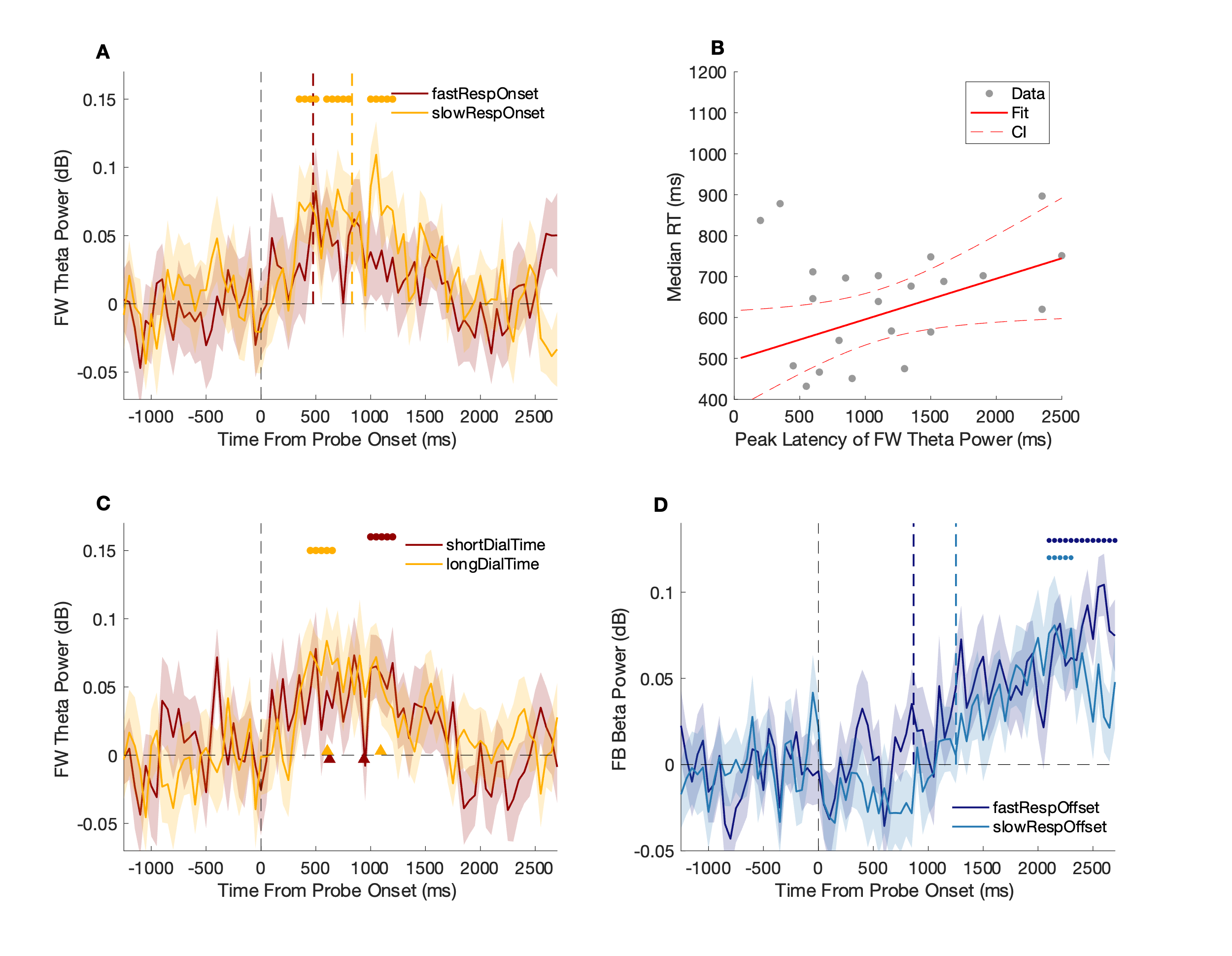

***Figure S6. Modest links between wave power and response times during no-precue trials in Experiment 2.*** *(A) Neither the amplitude nor the latency of FW theta waves varied with participants’ response onset times (median split). (B) We observed a modest but robust correlation between peak FW theta latency and individual differences in response times (r^2^ = 0.136,* *p = 0.0391). (C) Neither the amplitude nor the latency of FW theta waves varied with participants’ response duration (dial time; median split). Neither the amplitude nor the latency of FB beta waves varied with response offset times (median split). Shaded regions in each plot depict ±1 S.E.M. The vertical dashed line at time 0 in (A), (C), and (D) depict probe onset. Colored vertical dashed lines in (A), (C), and (D) depict median response onset times (fast vs. slow), response durations (short vs. long), and response offset times (fast vs. slow).*
